## Supplementary Figures and legends for "Topological stress triggers persistent DNA lesions in ribosomal DNA with ensuing formation of PML-nucleolar compartment"

Supplementary Figure 1

A

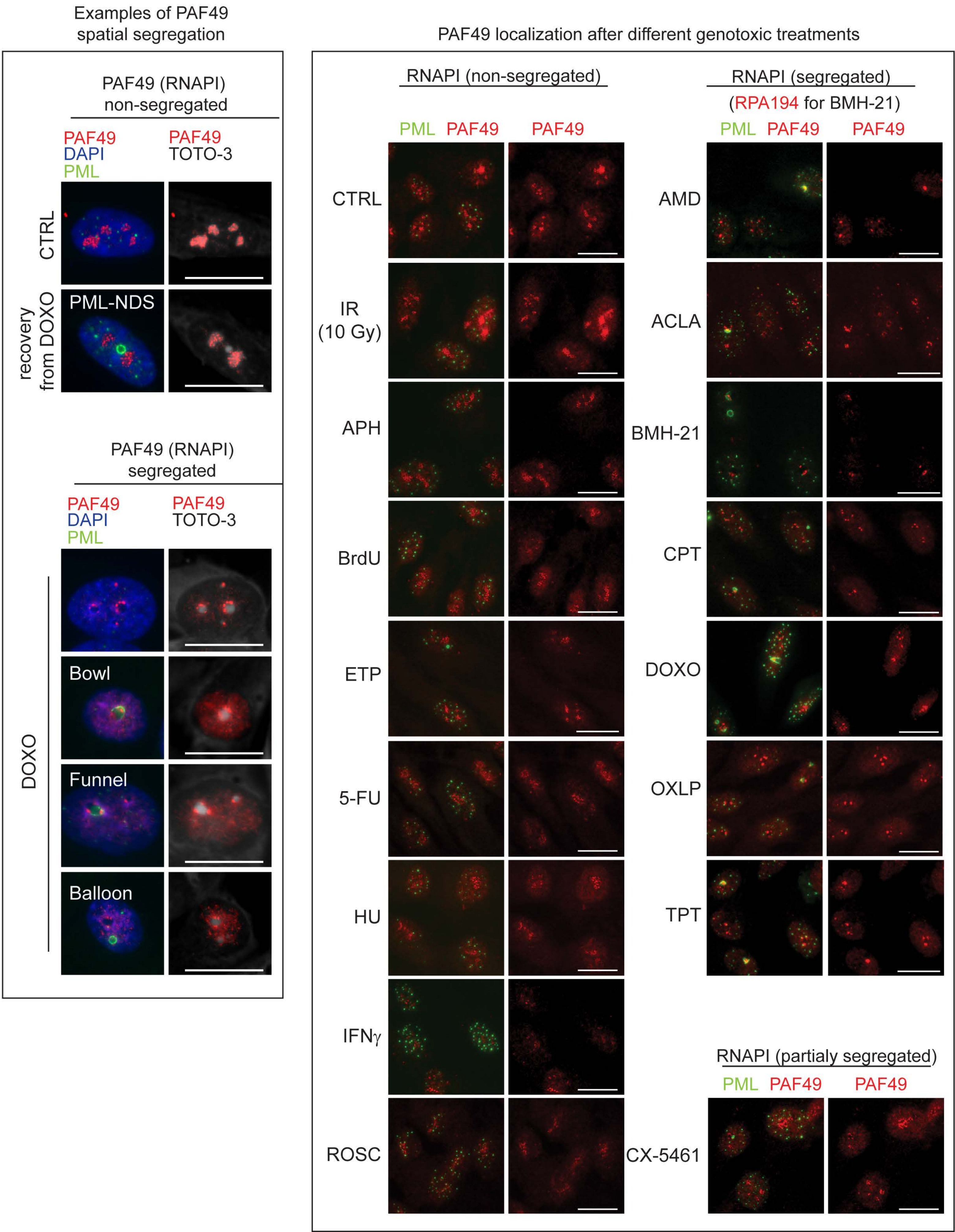

Supplementary Figure 1 (cont.)

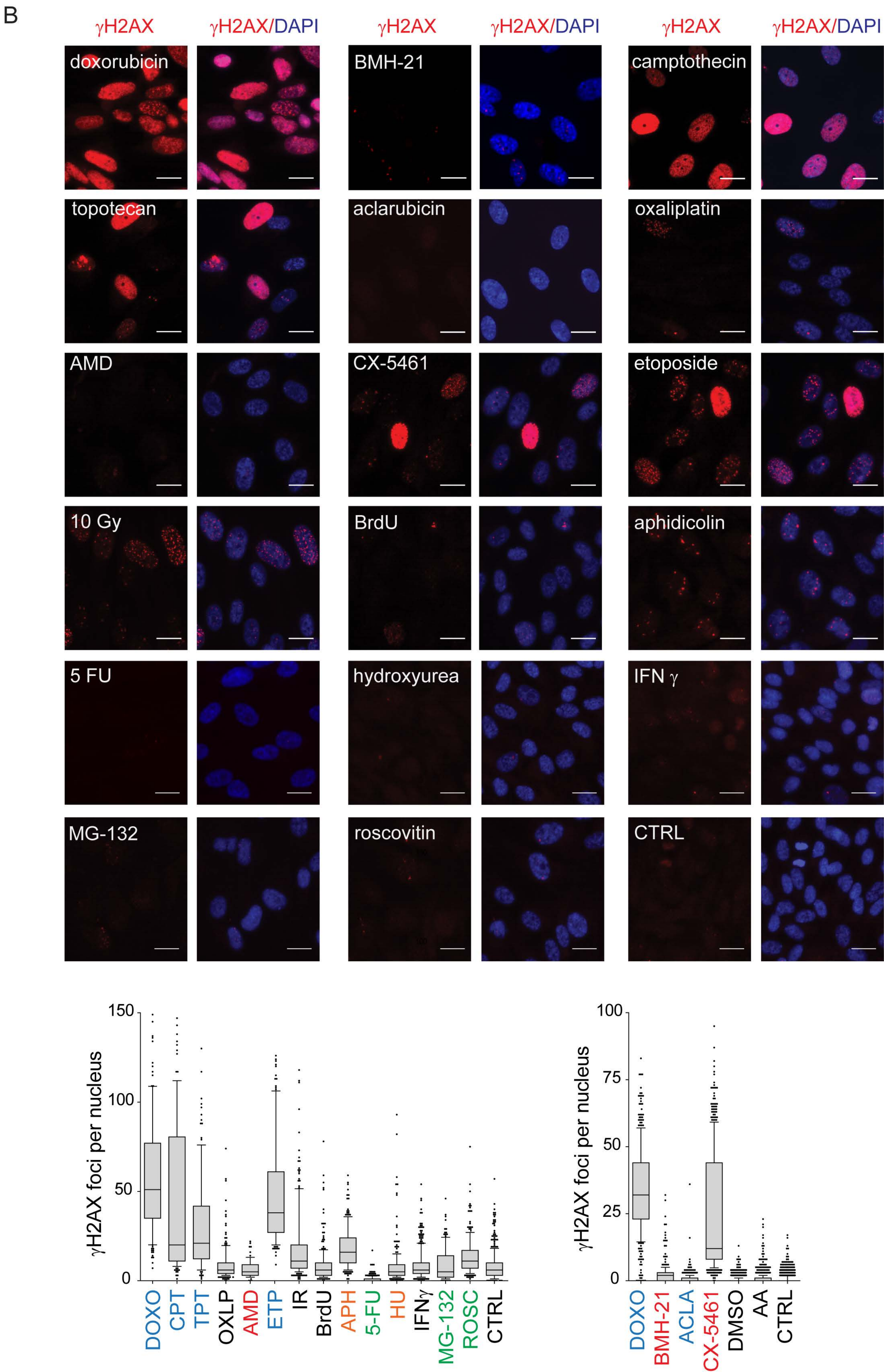

Supplementary Figure 1 (cont.)

C

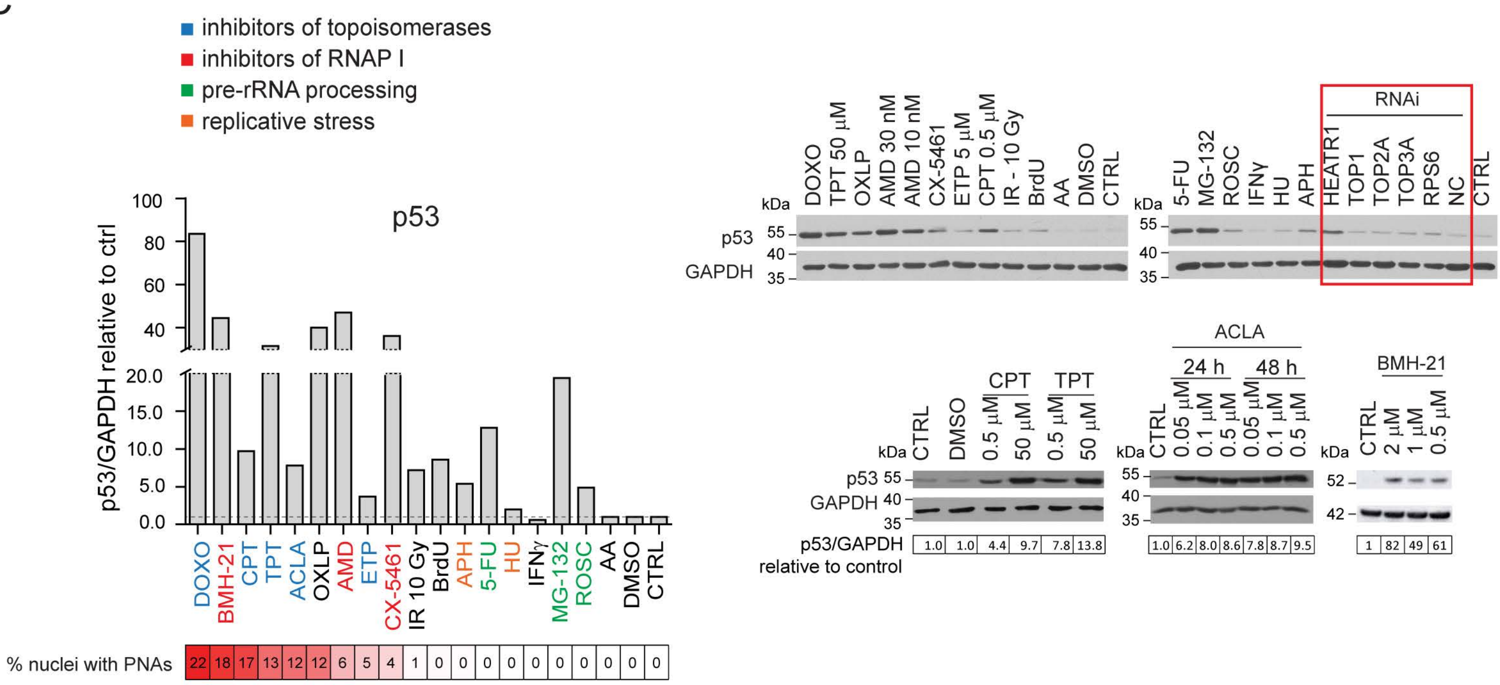


D

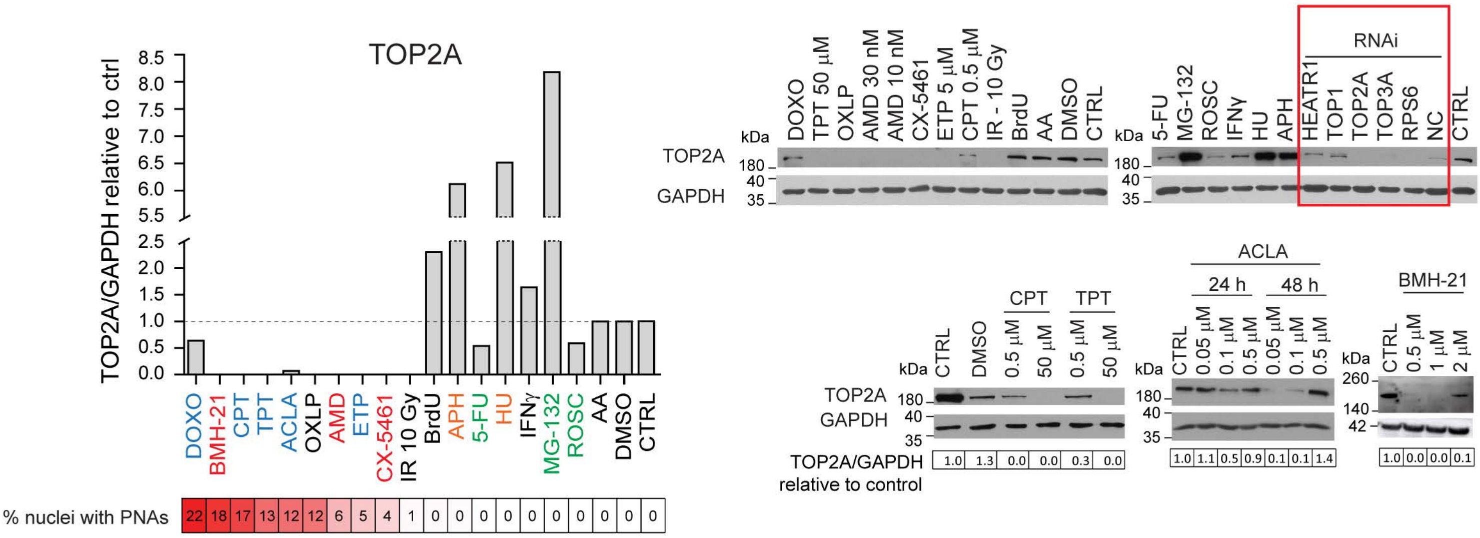


E

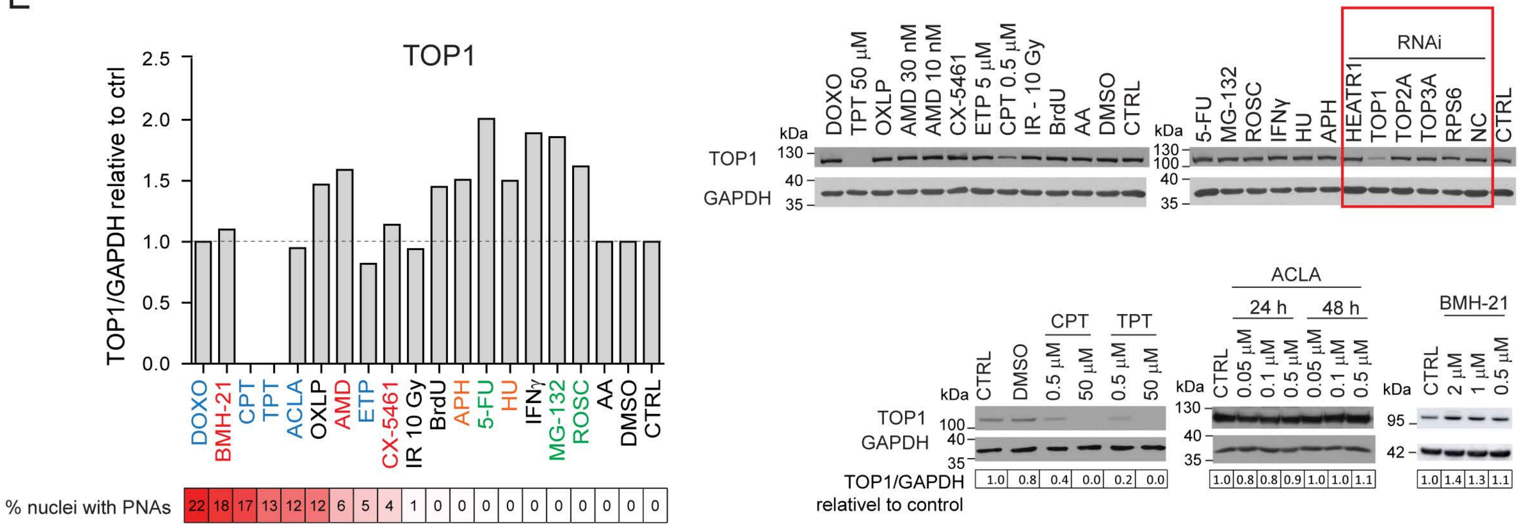

Supplementary Figure 1 (cont.)

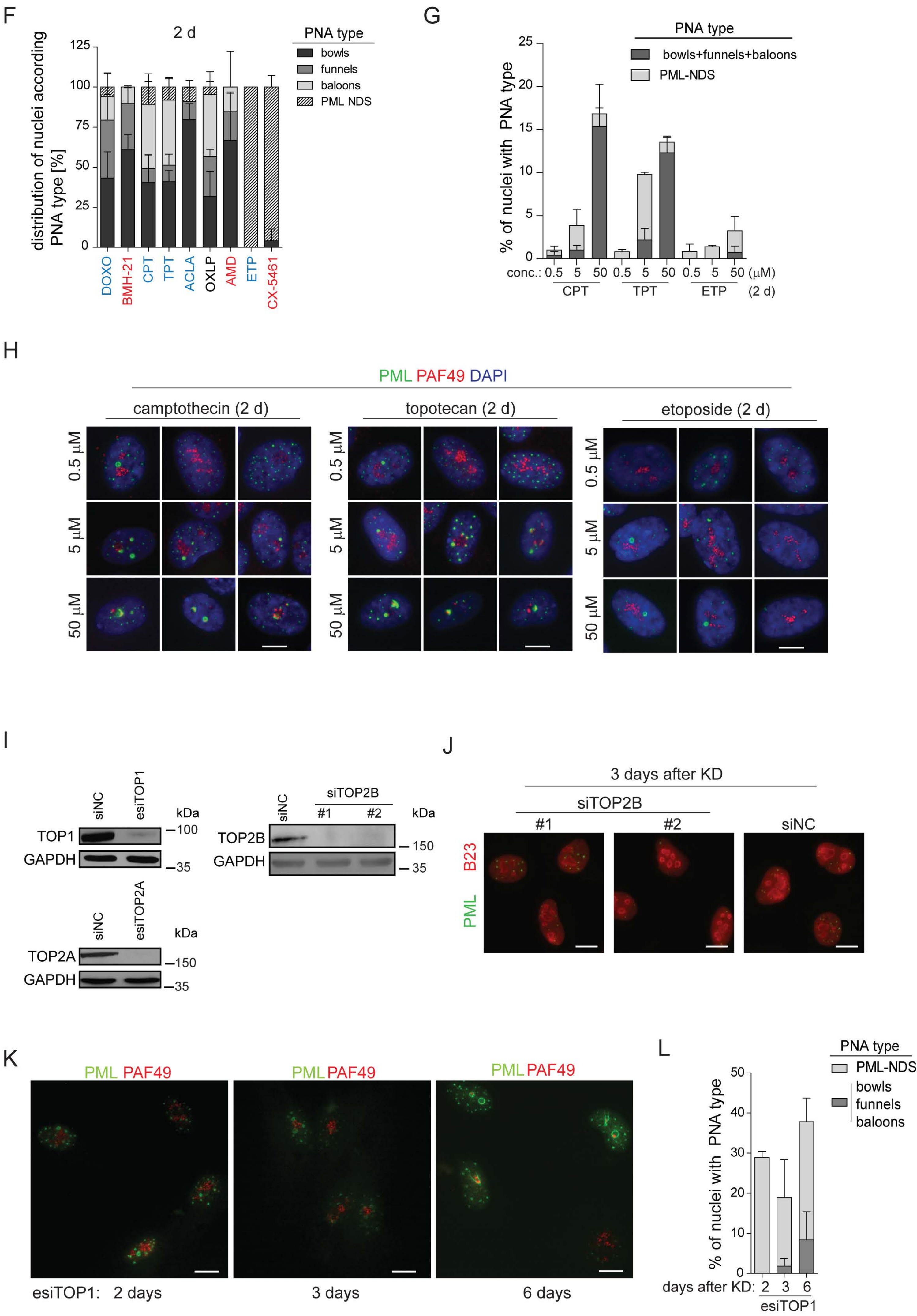

Supplementary Figure 1 (cont.)

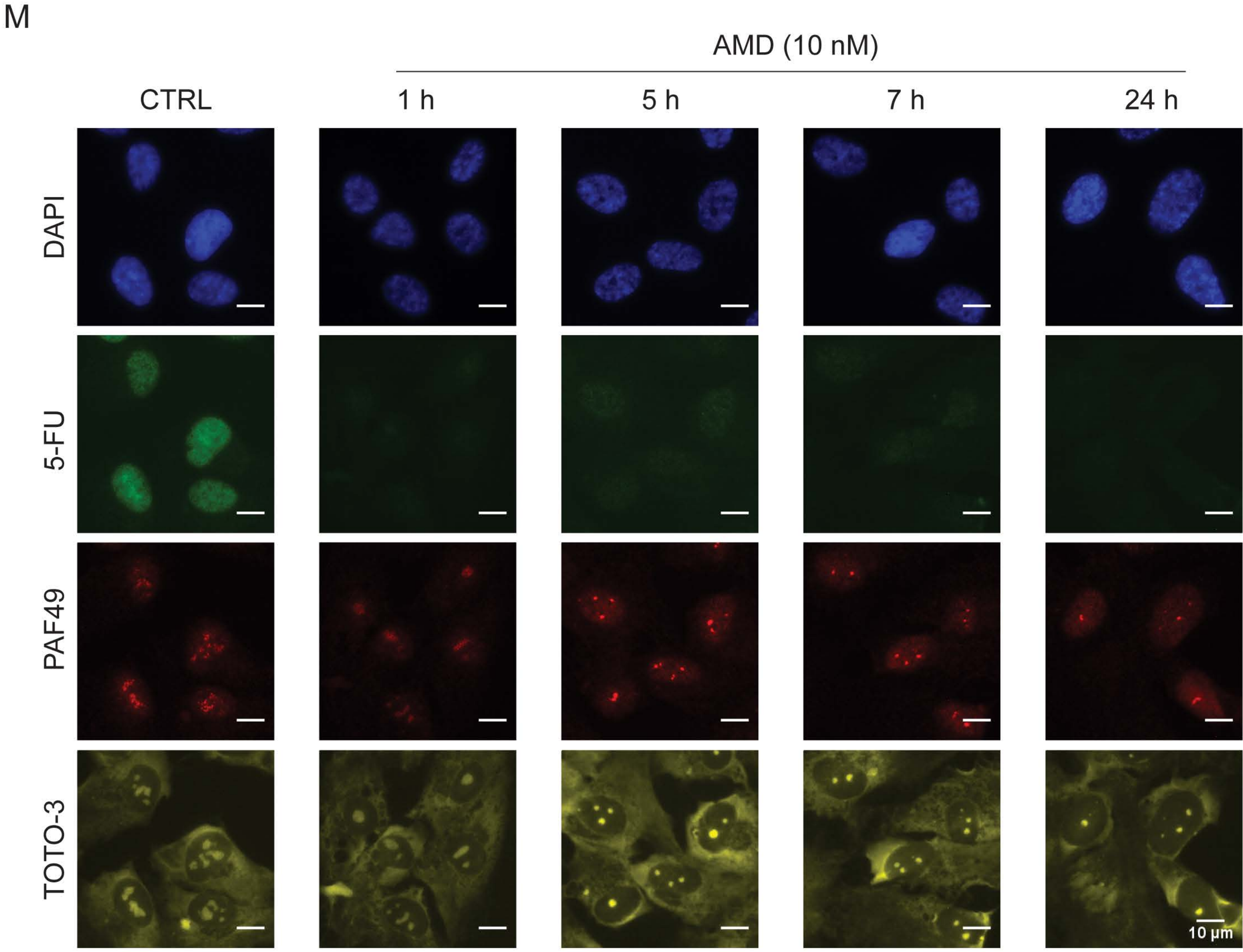


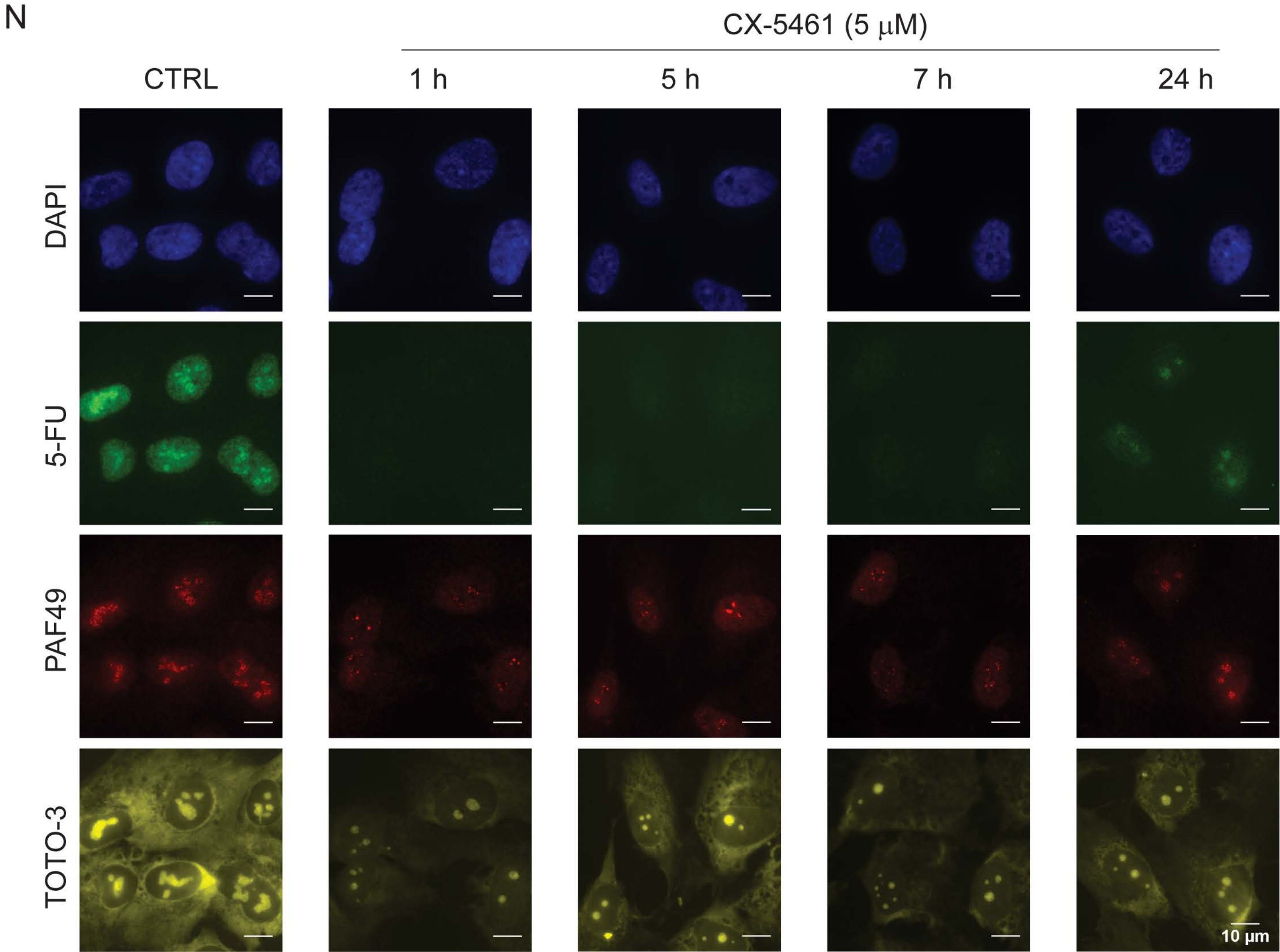

### Supplementary Figure 2

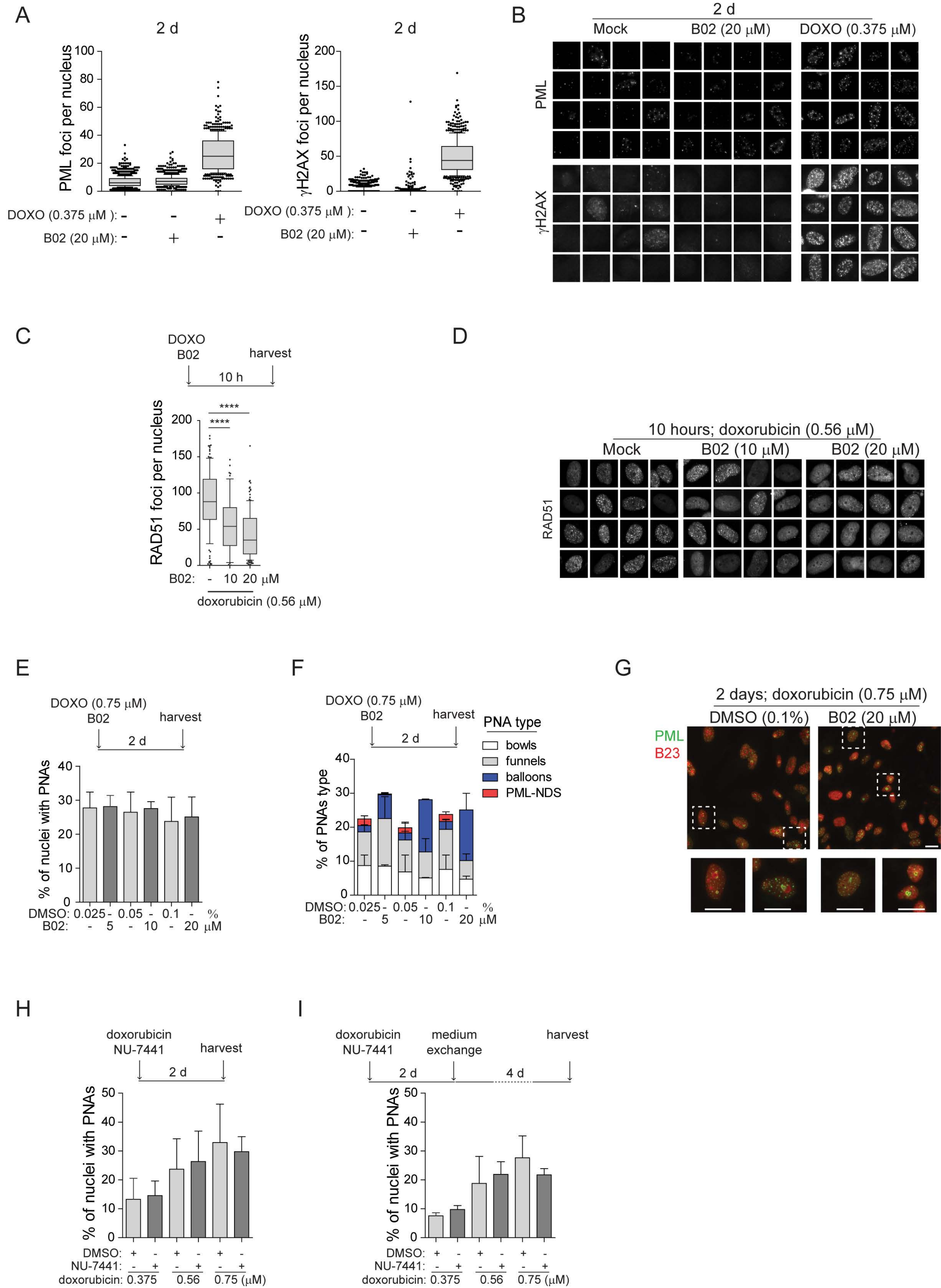

Supplementary Figure 2 (cont.)

J

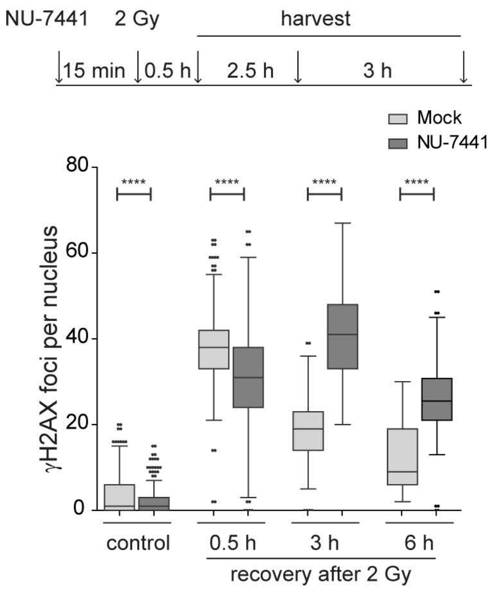


K

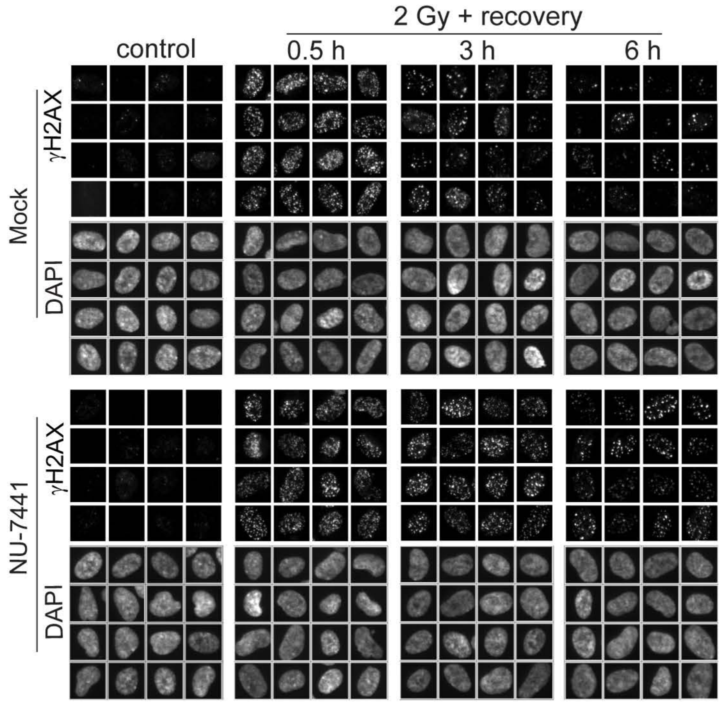


L

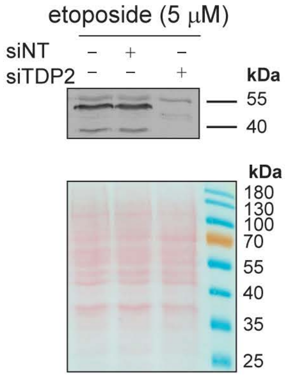


M

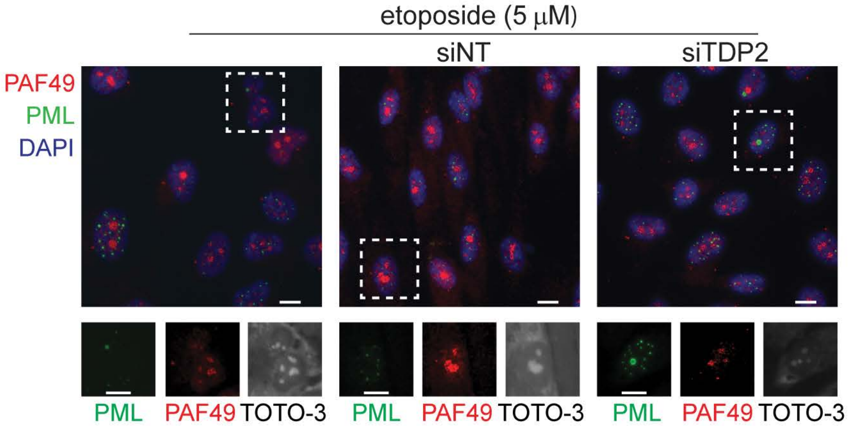


N

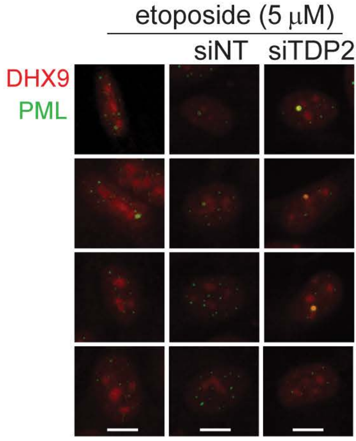

Supplementary Figure 3

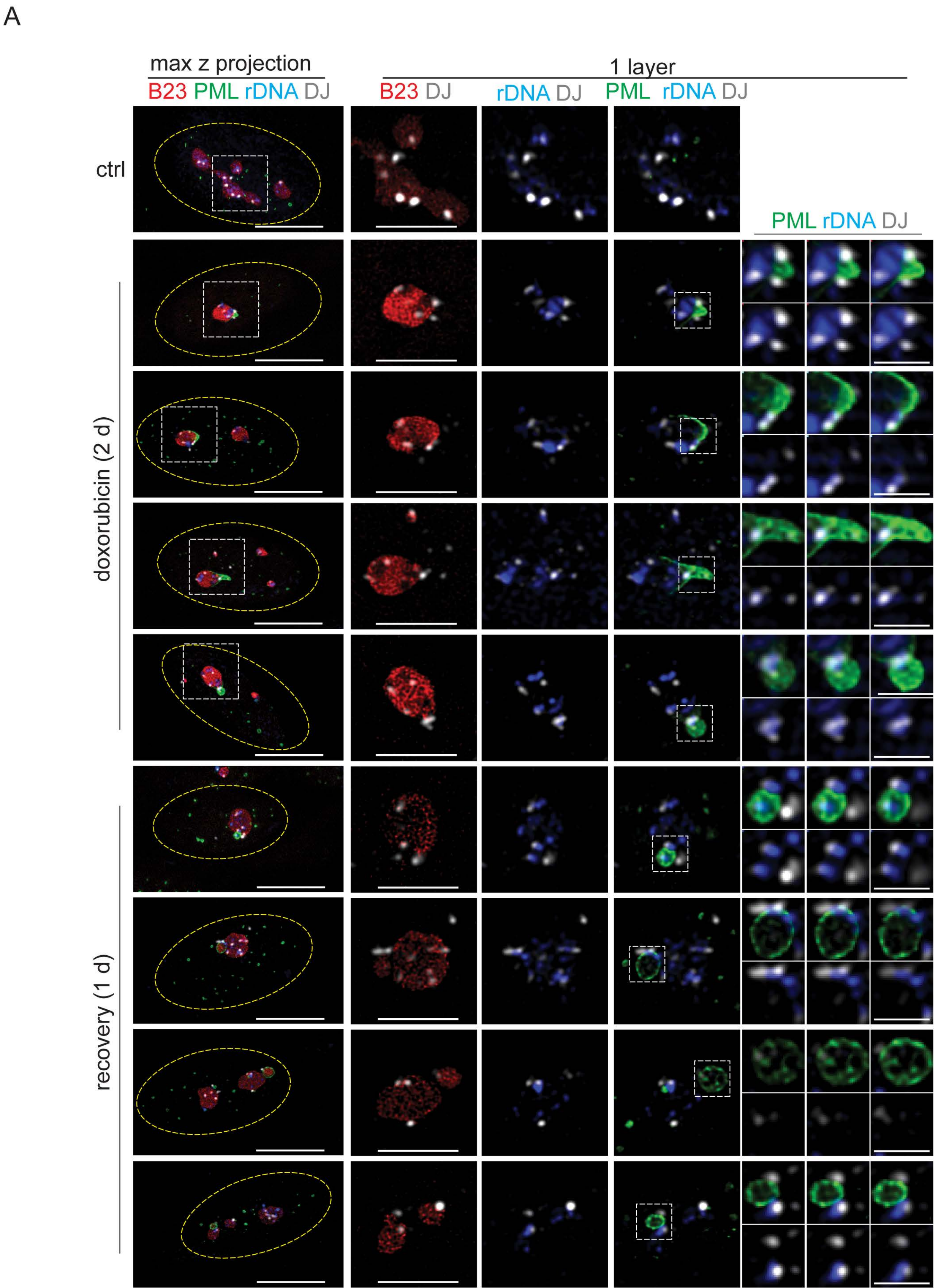

Supplementary Figure 3 (cont.)

B

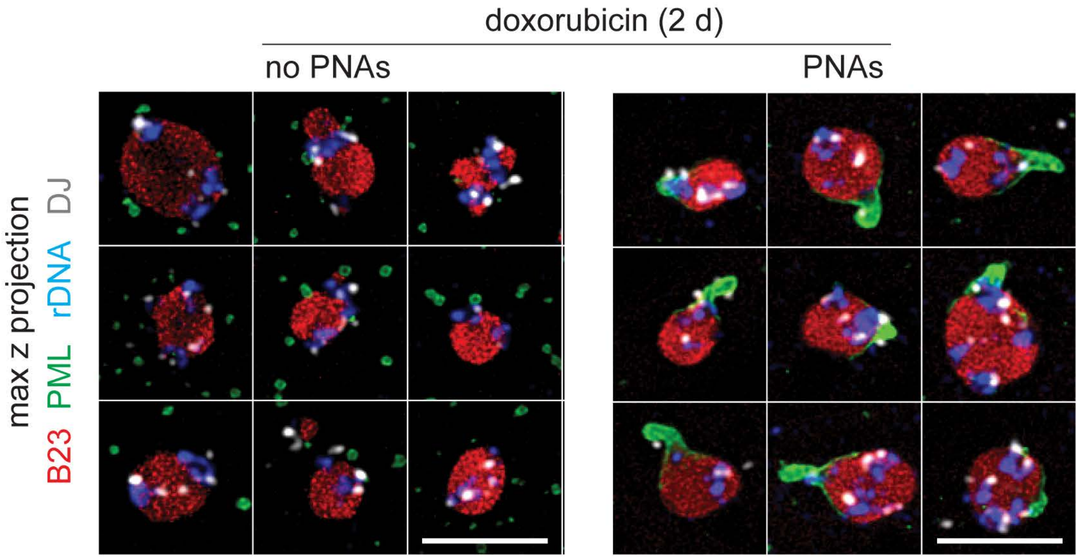


C

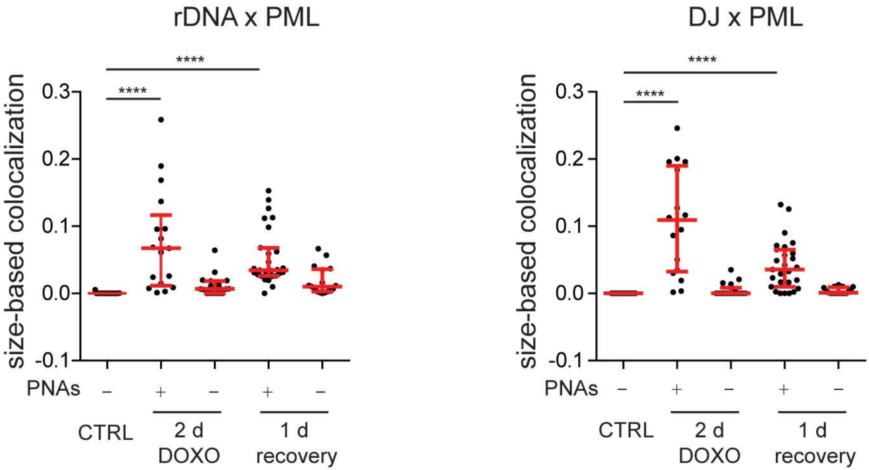

Supplementary Figure 3 (cont.)

D

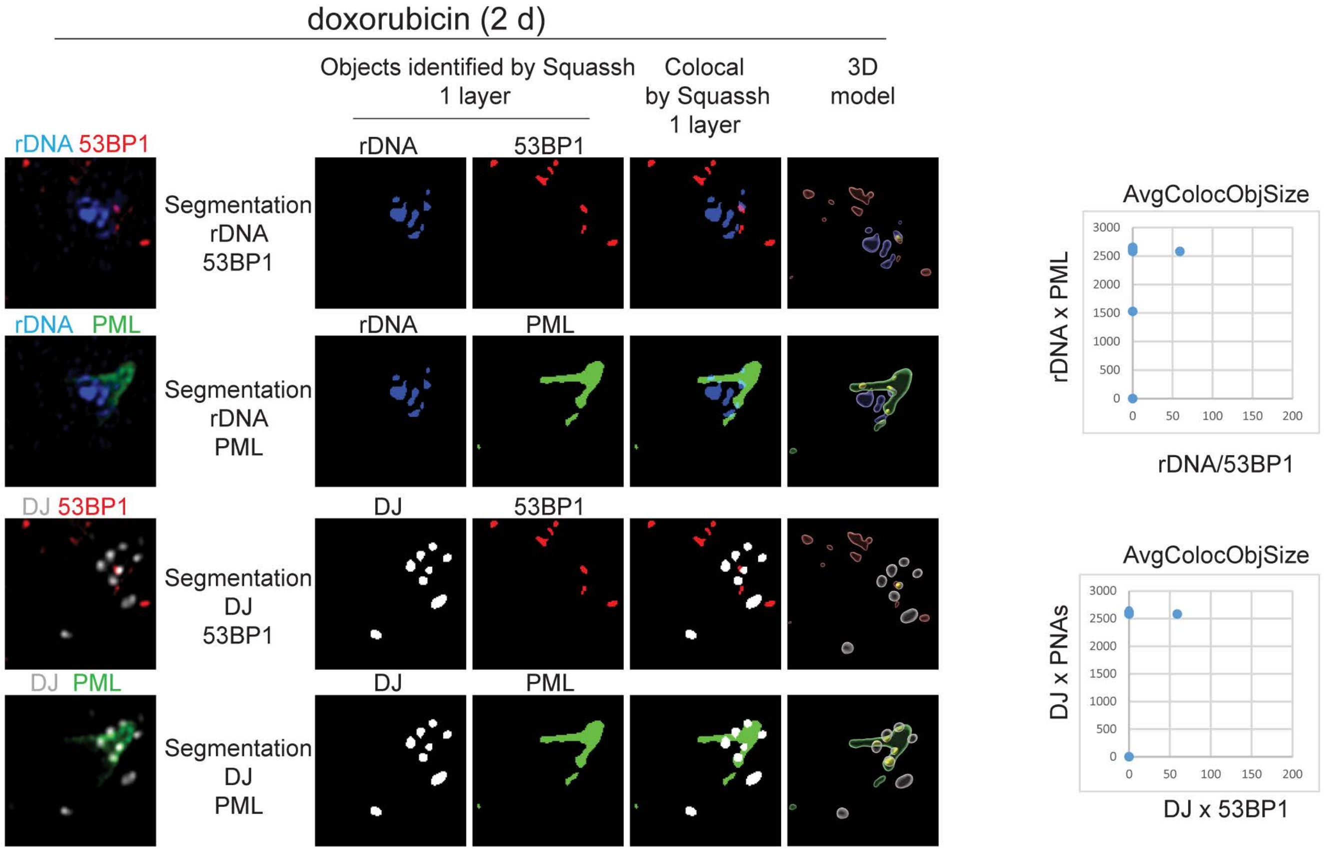


E

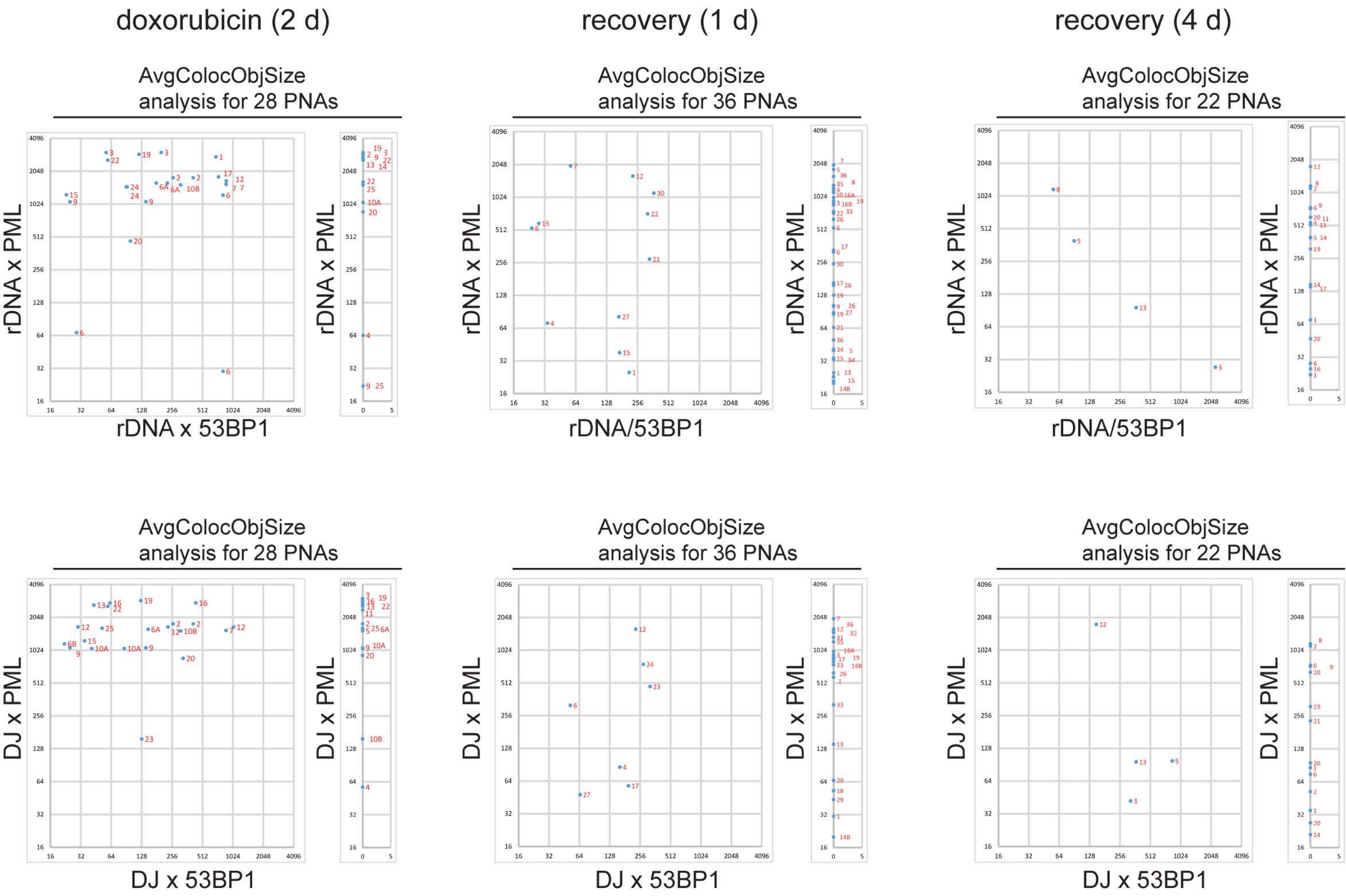

Supplementary Figure 4

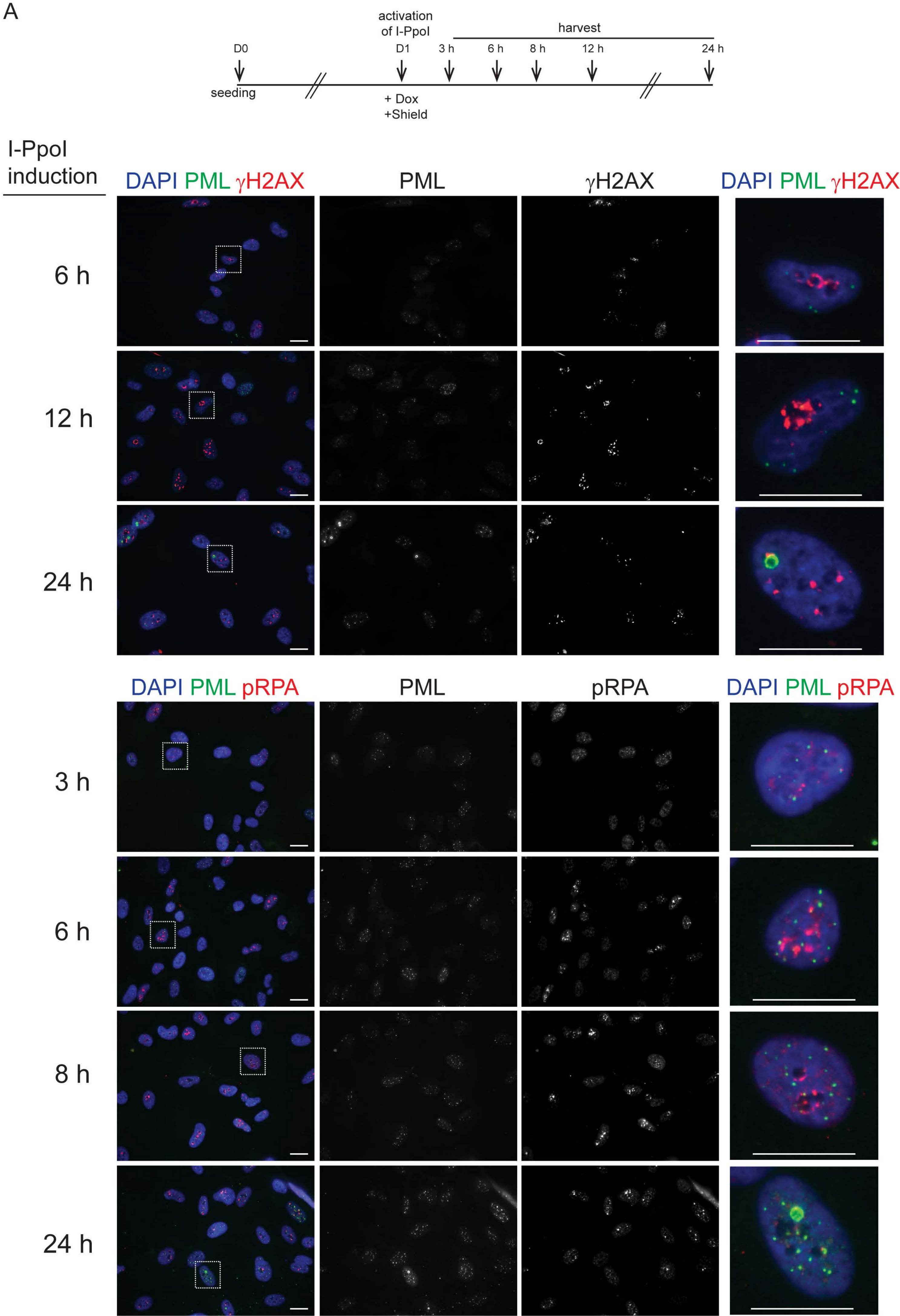

Supplementary Figure 4 (cont.)

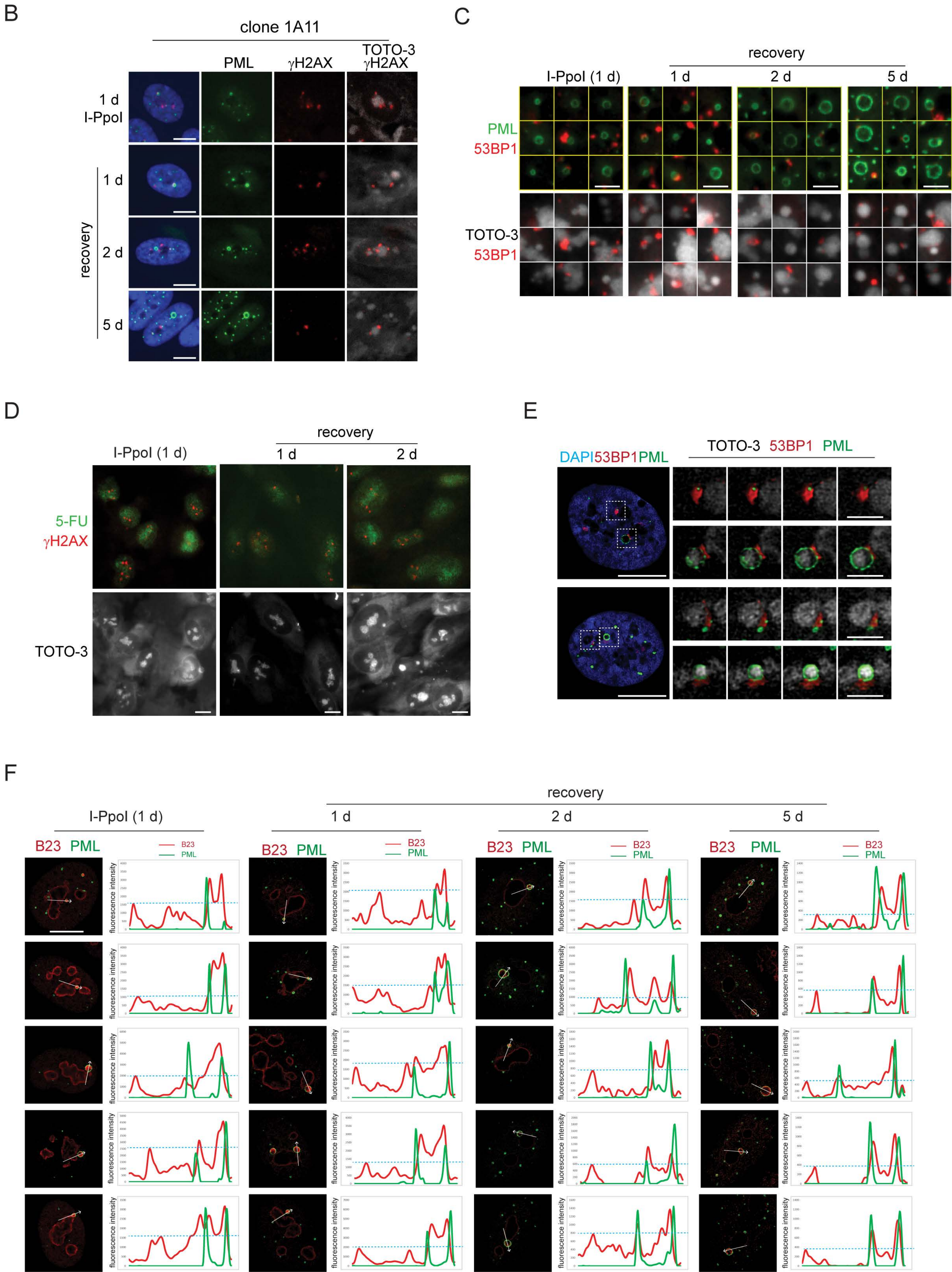

Supplementary Figure 4 (cont.)

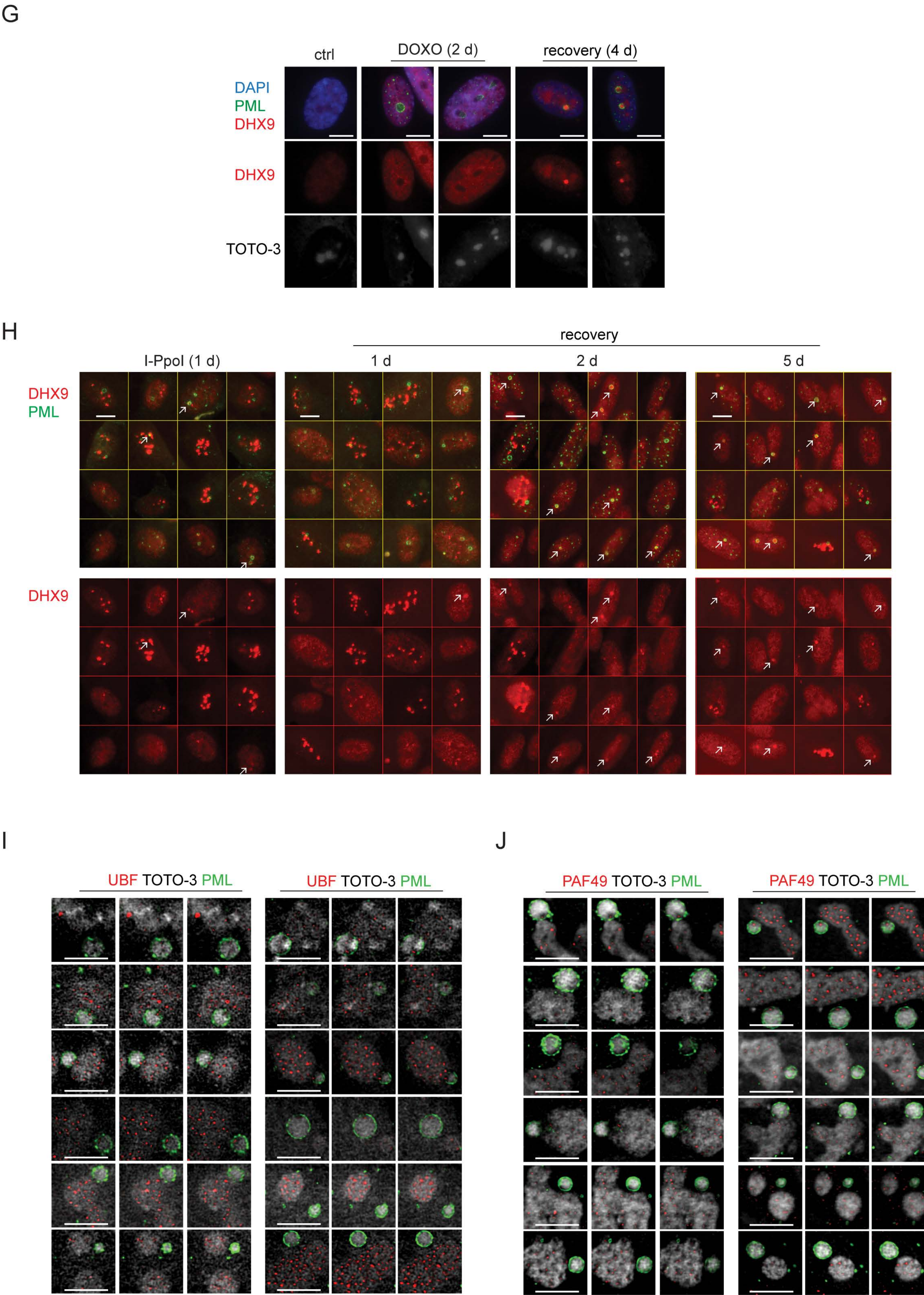

Supplementary Figure 4 (cont.)

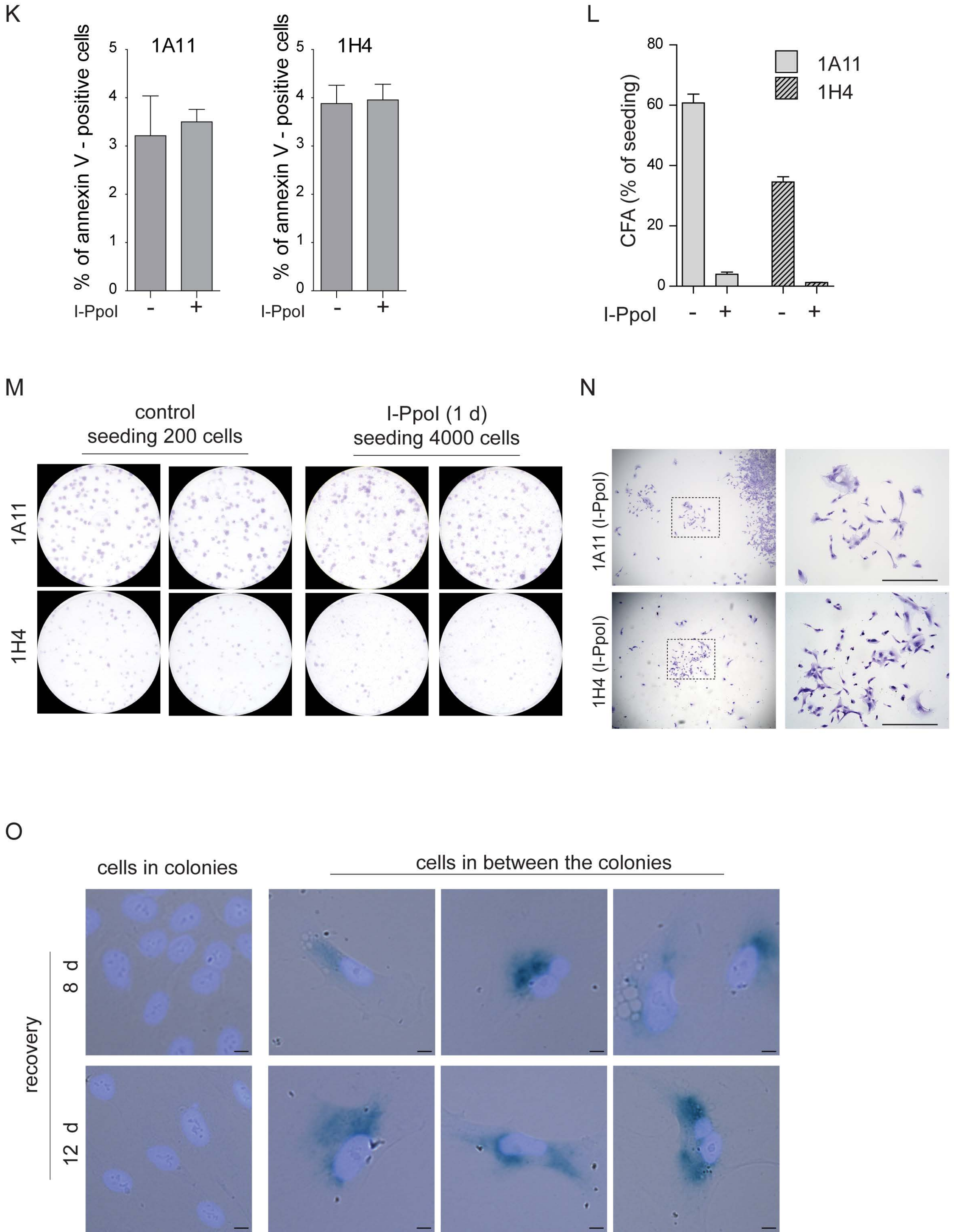

Supplementary Figure 5

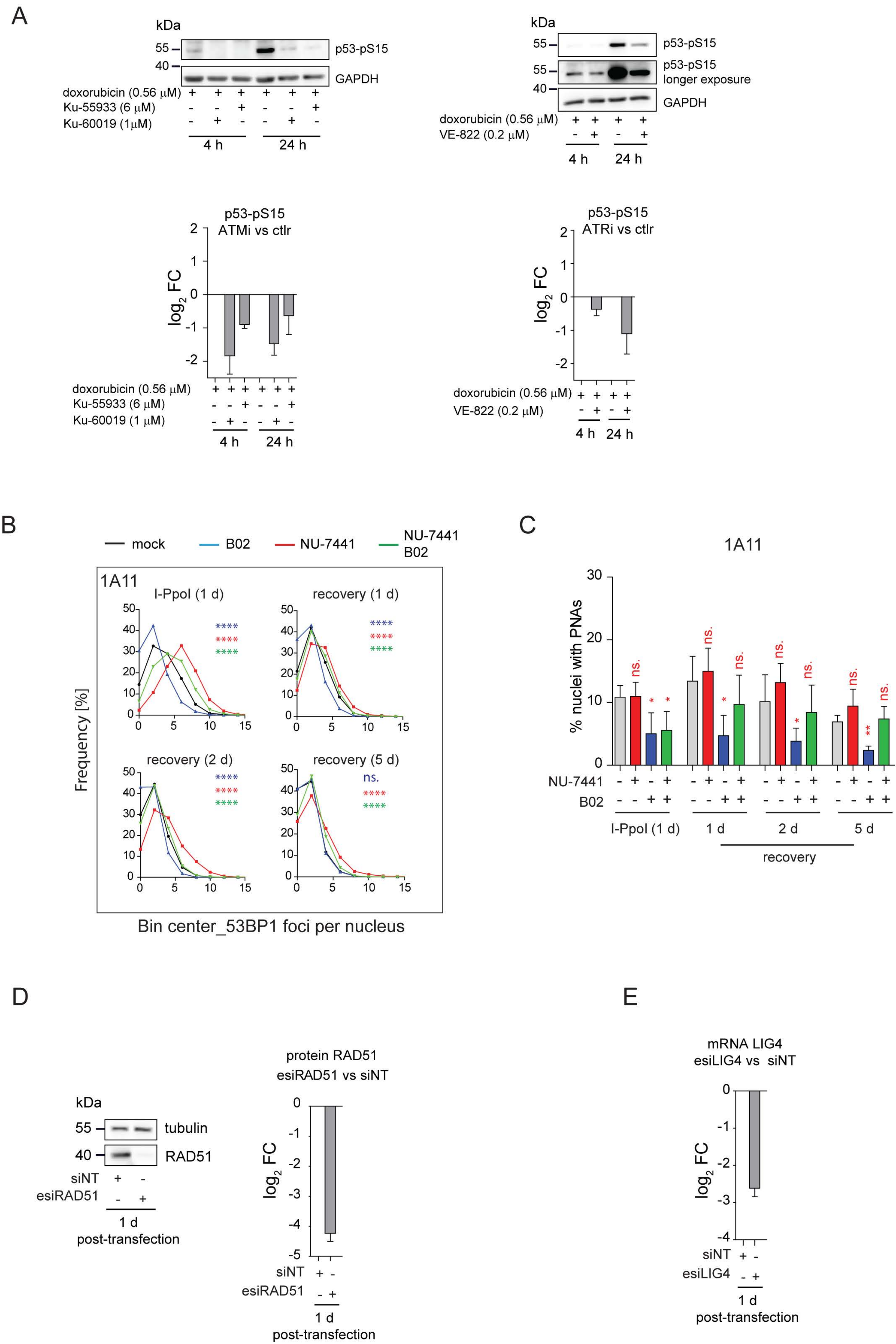

Supplementary Figure 5 (cont.)

F

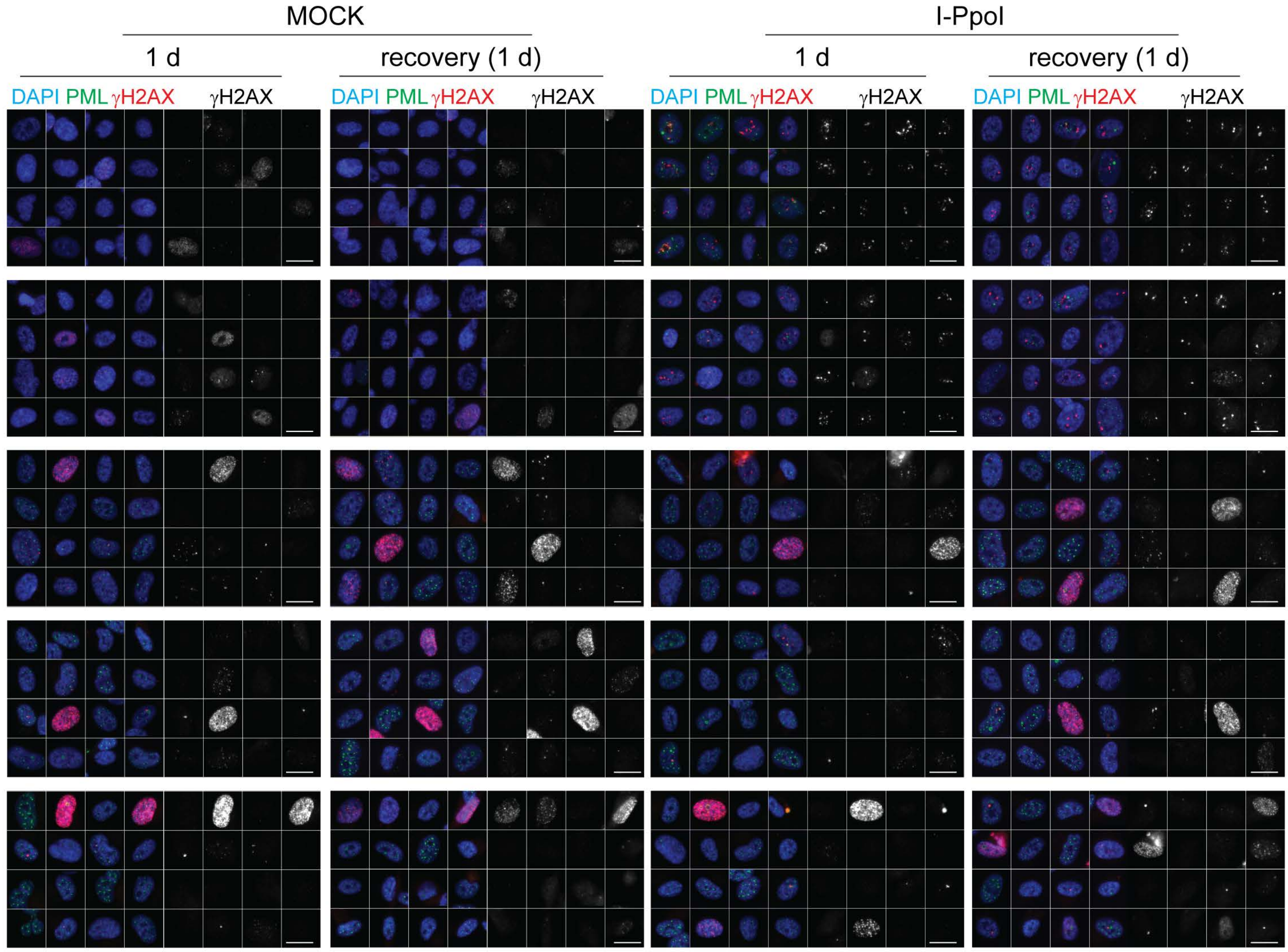


G

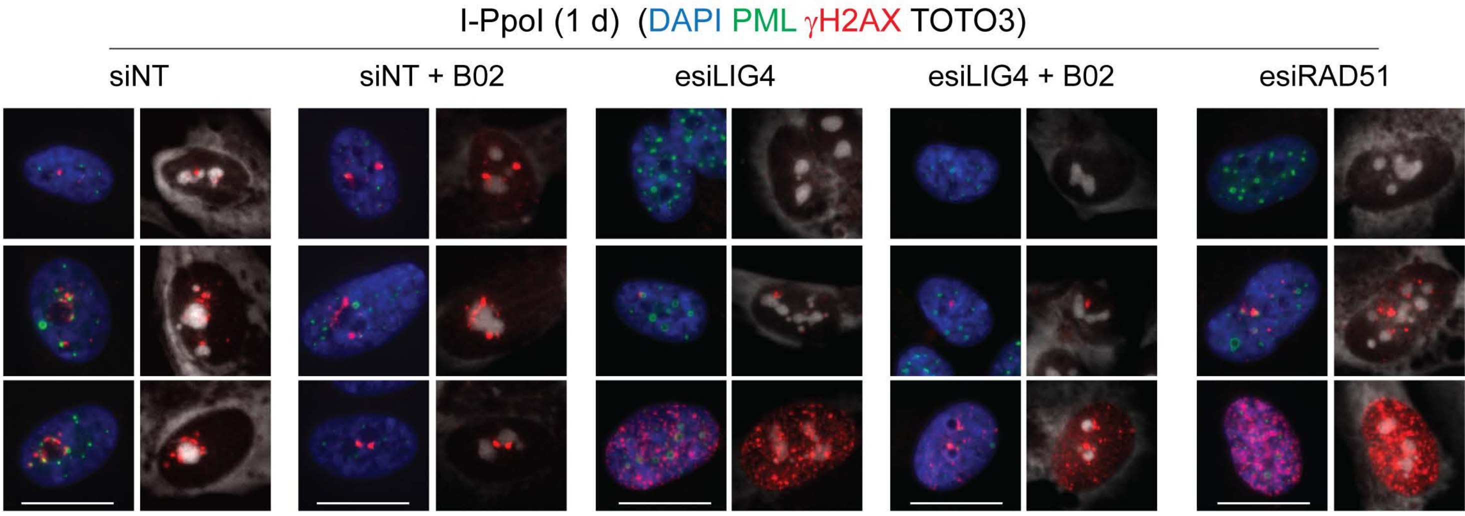


H

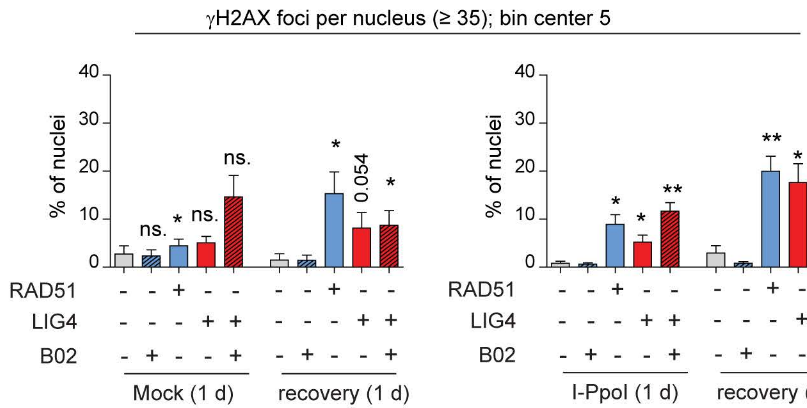


I

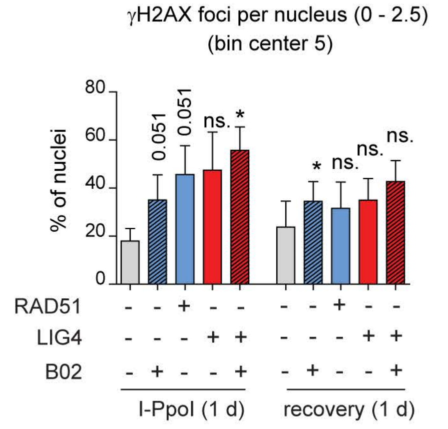

**Supplementary Figure 1. RNAPI segregation, γH2AX signal and the levels of p53, TOP1, and TOP2A after selected treatments; topoisomerase downregulation and RNAPI segregation after esiTOP1; RNAPI inhibition after AMD or CX-5461.** RPE-1$^{hTERT}$ cells were treated with selected treatments for 48 hours. (**A**) The segregated and non-segregated pattern of RNAPI subunits PAF49 (red) and different PNA subtypes (PML; green) are shown in the left panel. Nuclei were counter-stained with DAPI (blue) and nucleoli with TOTO-3 (white). In the right panel, the activity of RNAPI, evaluated as spatial segregation of RNAPI subunits PAF49 or RPA194 (BMH-21), and the presence of PNAs were visualized by wide-field immunofluorescence microscopy of PAF49/RPA194 (red) and PML (green). Scale bar, 20 µm. (**B**) The presence of DSBs, demonstrated as phosphorylation of histone H2AX on serine 139 (γH2AX), was visualized by indirect immunofluorescence using the anti-γH2AX antibody (red), while the nuclei were counter-stained with DAPI (blue). The γH2AX count per nucleus was analyzed using ScanR software and is presented using a Whiskers plot (box: 10 – 90 percentiles; black line: median). Scale bar, 20 µm. (**C**, **D**, and **E**) RPE-1$^{hTERT}$ cells were treated with selected treatments for 48 hours and the levels of p53, TOP2A, and TOP1 were detected by western blotting. The charts on the left show relative levels of p53, TOP2A, and TOP1, respectively, related to the loading control. The protein levels of untreated cells or cells treated with drug solvents (acetic acid, DMSO) were set as one. The red, one-row tables below the charts show the percentage of cells containing PNAs. (**F**) RPE-1$^{hTERT}$ cells were treated with doxo, BMH-21, CPT, TPT, acla, OXLP, AMD, CX-5461, and ETP for 48 h (for the used concentration, see Supplementary Table 1). The distribution of nuclei containing either bowls, funnels, balloons, or PML-NDS was evaluated for three independent experiments. Results are presented as a mean ± s.d. (**G**) RPE-1$^{hTERT}$ cells were treated with three different concentrations of CPT, TPT, or ETP for 48 hours, and percentages of cells containing either bowls/funnels/balloons or PML-NDS were evaluated for three independent experiments. Results are presented as a mean ± s.d. (**H**) The localization of PAF49 (a subunit of RNAPI; the level of segregation indicates the RNAPI activity) after treatments described in (G) is shown. The PAF49 (red), and PML (green) were detected by indirect immunofluorescence and wide-field microscopy. The nuclei were co-stained by DAPI (blue). Scale bar, 10 µm. (**I**) Immunoblotting of lysates extracted from cells after transfection either with non-targeting siRNA or with esiRNA/siRNA targeting individual topoisomerases. The membranes were probed with indicated anti-Top antibodies to prove their efficient downregulation. GAPDH was used as a loading control. (**J**) Illustrative images of RPE-1$^{hTERT}$ cells in which TOP2B was downregulated by two different siRNAs are shown. PML (green) and B23 (red; a nucleolus marker) were detected by indirect IF and wide-field microscopy. Scale 10 µm. (**K**) Illustrative images of RPE-1$^{hTERT}$ cells in which TOP1 was downregulated by esiRNA transfection show time-dependent changes in the PML (green) and PAF49 (red) patterns. While 2 days after the transfection PAF49 was mostly non-segregated and PML-NDS were the most frequent PNAs appeared, 6 days after the transfection PAF49 was mostly segregated, and PML-NDS were largely substituted by PML forks and circles. Scale bar, 10 µm. (**L**) The quantification of (K). The percentages of cells containing either bowls/funnels/balloons or PML-NDS were evaluated.

Results are presented as a mean ± s.d. (**M, N**) RPE-1$^{hTERT}$ cells were treated with AMD or CX-5461 for indicated time points. At each time point, the cells were incubated with 5-FUrd for 30 min. The immunofluorescence detection of 5-FUrd (green), representing newly synthesized nucleolar RNA, and localization/segregation of RNAPI subunit PAF49 (red) was performed. The nuclei and nucleoli were stained with DAPI (blue) and TOTO-3 (yellow), respectively. Scale bars, 10 μm.

**Supplementary Figure 2. The inhibition of specific DNA repair pathways augmented the formation of PNAs.** (**A**) RPE-1$^{hTERT}$ cells were exposed to 20 μM B02, the corresponding concentration of DMSO (mock), and 0.375 μM doxorubicin for 48 h to control whether treatment with B02 can induce the PNAs or DSBs. PML and γH2AX were detected by indirect IF and high-content microscopy unit (ScanR). The PML or γH2AX count per nucleus is shown as Whiskers (box: 10 – 90 percentiles; black line: median). (**B**) The galleries of nuclei with PML or γH2AX foci after treatment described in (A) are shown. (**C**) RPE-1$^{hTERT}$ cells were treated with 0.56 μM doxorubicin and 10 μM and 20 μM B02 to confirm the inhibition of RAD51 filament formation by B02. RAD51 was detected using indirect IF and ScanR. The RAD51 count per nucleus is shown as Whiskers (box: 10 – 90 percentiles; black line: median). Only G2 cells discriminated by DNA content were analyzed. (**D**) The gallery of nuclei gated according to DAPI intensity for the G2 population with RAD51 foci after treatment described in (C) is shown. (**E**) RPE-1$^{hTERT}$ cells were exposed to 0.75 μM doxorubicin with three concentrations of B02 (5, 10, and 20 μM), or corresponding concentrations of DMSO. The plot represents the percentage of nuclei with PNAs after a 2-day-long treatment. (**F**) The plot represents the distribution of individual types of PNAs (%) after the treatments described in (E). (**G**) Representative cell from (E) after 2-day-long treatment with 0.75 μM doxorubicin combined with 20 μM B02 or 0.1% DMSO (mock). (**H**) RPE-1$^{hTERT}$ cells were exposed to three concentrations of doxorubicin (0.375 μM, 0.56 μM, and 0.75 μM) in combination with 1 μM NU-7441 (inhibitor of DNA PK and NHEJ) or corresponding concentration of DMSO (mock). The plot represents the percentage of nuclei with PNAs after a 2-day-long treatment. (**I**) RPE-1$^{hTERT}$ cells were exposed to three concentrations of doxorubicin (0.375 μM, 0.56 μM, and 0.75 μM), combined with 1 μM Nu7441 (inhibitor of DNA PK and NHEJ) or DMSO. The plot represents the percentage of nuclei with PNAs after a 2-day-long treatment and 4-day-long recovery. (**J**) RPE-1$^{hTERT}$ cells were exposed to ionizing radiation (2 Gy) in the presence or absence of 1 μM Nu7441 (added 15 min before irradiation). The presence of γH2AX foci (a marker of DSB s) was analyzed 0.5, 3, and 6 hours after ionizing radiation by indirect immunofluorescence and ScanR. Quantification of γH2AX count per nucleus is presented as a whiskers box with a median. Only G1 cells discriminated by DNA content were analyzed. (**K**) The gallery of nuclei (co-stained with DAPI) with γH2AX foci treated and analyzed as described in (J) is shown. (**L**) RPE-1$^{hTERT}$ cells were transfected with siRNA targeting TDP2 or non-targeting siRNA as control. 2 days after the transfection, the 5 μM etoposide was added for an additional 2 days. Then, the cells were harvested, and the level of TDP2 protein was examined. The Ponceau-S staining is shown as a control. (**M**) RPE-1$^{hTERT}$ cells were treated as described in (L), and the PML (green) and PAF49 (subunit of RNAPI; red) were detected by indirect IF and wide-field microscopy to evaluate the activity of RNAPI marked as PAF49 segregation and correlation with PNAs type occurrence. The nuclei and nucleoli were co-stained by DAPI and TOTO-3, respectively. (**N**) After the treatment described in (L), the PML and DHX9 were detected by indirect IF and wide-field microscopy to prove the accumulation of DHX9 in PML-NDS. In all experiments, at least three biological replicates were evaluated. Results are presented as a mean ± s.d. Student's t-test was used for statistical evaluation. Asterisks indicate the following: ****P < 0.0001, ***P < 0.001, **P < 0.01, *P < 0.05. Scale bars, 20 μm (G) and 10 μm (M and N).

**Supplementary Figure 3. PNAs envelop rDNA and DJ loci containing rDNA DSB after doxorubicin treatment**. (**A**) RPE-$1^{hTERT}$ cells were treated with 0.75 μM doxorubicin for 2 days and recovered from the treatment for 1 day. The proliferating cells were used as a control. The localization of rDNA, DJ, PML, and B23 was analyzed using immuno-FISH staining and confocal microscopy to find the correlation between nucleolar caps (individual NORs) and PNAs. The sections of several nucleoli and the associated PNAs are presented in detail: rDNA (blue), DJ (white), PML (green), and B23 (red). (**B**) The gallery of nucleoli with and without PNAs after treatment, staining, and capturing described in (A) is presented. (**C**) The extent of PML-rDNA and PML-DJ size-based colocalization calculated for individual nucleoli of treated and untreated cells with respect to the presence of PNAs is shown as a scatter plot. The median with an interquartile range is shown. The colocalization was calculated using Fiji/Mosaic/Squassh plugin. The number of analyzed nucleoli in each group was: ctrl (n=19); 2 days + PNAs (n=17); 2 days without PNAs (n=18); 1-day-long recovery + PNAs (n=27); 1-day-long recovery without PNAs (n=17). (**D**) The image of representative nucleolus with funnel-like PNAs is used to explain the principle of analysis used to identify whether PNAs co-localize with rDNA/DJ with DSB. The colocalization between rDNA/DJ and PML and rDNA/DJ and 53BP1 was done using Fiji/Mosaic/Squassh plugin. The original image, the segmentation, the obtained co-localization between individual objects, and 3D model (done in Imaris) are shown. As for both segmentations (rDNA–53BP1×rDNA–PML; similarly for DJ–53BP1×DJ–PML) the same parameters were used, and each object got the same number, the extent of colocalization with 53BP1 or PML with the same rDNA/DJ object could be defined. The presented scatter plot expresses the size of co-localization of individual rDNA/DJ objects with 53BP1 (x-axis) and PML (y-axis). (**E**) The size of colocalization of individual rDNA/DJ objects with PML and 53BP1 obtained for all analyzed nucleoli is shown in the x/y scatter plot. Scale bars, 10 μm (A-nuclei), 5 μm (A, B-nucleoli), and 2 μm (A-PNAs).

**Supplementary Figure 4. The DNA damage introduced into the rDNA locus by endonuclease I-PpoI induces PML-NDS and cellular senescence**. (**A**) The I-PpoI was activated 24 hours after RPE-$1^{hTERT}$-I-PpoI (1H4) seeding using doxycycline and Shield. The cells were harvested 3, 6, 8, 12, and 24 hours after I-PpoI induction. The PNAs (PML; green) and DSBs (γH2AX or RPA2pS33; red) were detected by indirect IF and ScanR. The DNA was stained by DAPI (blue). (B – F and H – J) The expression of I-PpoI was induced in RPE-1$^{hTERT}$ -I-PpoI cells for 24 h, then the medium was exchanged, and different parameters were observed during the recovery phase. (**B**) The representative images obtained by indirect IF and wide-field microscopy show the localization of γH2AX (a marker of DSB; red) and PML (green) and their relationship with the nucleolus. The nuclei and nucleoli were co-stained with DAPI (blue), and TOTO-3 (white), respectively. (**C**) The gallery of PNAs (PML-NDS) indicates their relationship with DSB and nucleoli. Indirect IF and wide-field microscopy detected 53BP1 (a marker of DSB; red) and PML (a marker of PML-NDS; green). The nucleoli were stained by TOTO-3 (white). (**D**) The activity of RNAPI was detected by 5-FU by 30 min long incubation at different time points. 5-FU (green) represents newly synthesized nucleolar RNA and γH2AX (red) DSBs. The nucleoli were co-stained TOTO-3 (white). (**E**) The localization of DSB next to PML-NBs and PML-NDS is shown. The 53BP1 (a marker of DSB; red) and PML (a marker of PML-NBs and PML-NDS; green) were detected by indirect IF and confocal microscopy (Stellaris). The nuclei and nucleoli were stained with DAPI (blue) and TOTO-3 (white), respectively. (**F**) The accumulation of B23 in PML-NDS is presented by plot profiles representing the intensity of B23 (red) and PML (green) fluorescence. The images obtained by indirect IF and confocal microscopy (Stellaris) were analyzed. (**G**) The localization of DHX9 after exposure of RPE-1$^{hTERT}$ cells to 0.75 μM doxorubicin is shown after a 2-d treatment followed by a 4-d recovery. DHX9 (red) and PML (green) were detected by indirect IF and wide-field microscopy. The nuclei and nucleoli were stained with DAPI (blue) and TOTO-3 (white), respectively. (**H**) The gallery of nuclei represents the link between the DHX9 localization and PML-NDS after the I-PpoI insult and the recovery. The DHX9 (red) and PML (a marker of PML-NDS, green) were detected by indirect IF and wide-field microscopy. The arrows indicate PML-NDS with DHX9 inside. Note this localization of DHX9 is characteristic of doxorubicin-induced PML-NDS. (**I** and **J**) The representative images show the link between UBF (I) or PAF49 (J) and I-PpoI-induced PML-NDS. UBF (a marker of rDNA; red), PAF49 (a subunit of RNAPI; red), and PML (a marker of PML-NDS; green) were detected by indirect IF and confocal microscopy (Stellaris). The nucleoli were stained with TOTO-3 (white). Three sequential layers are presented. (**K**) The I-PpoI was activated for 24 h then cells were analyzed for marker of apoptosis, annexin-V using Fluorescence-Activated Cell Sorting (FACS). The percentage of annexin V-positive cells is shown as a mean from 2 biological replicates. (**L**) The ability to form colonies after treatment described in (K) was assessed using colony forming assay. The percentage of cells able to form a colony is shown as a mean from three independent biological replicates. (**M** and **N**) The representative plates from (L) and the morphology of cells stained by crystal violet and captured by ZEISS AxioZoom V.16 microscope are shown. (**O**) In cells treated as (K) the senescence-associated β-galactosidase activity was detected by colorimetric assay after 8 and 12 days of recovery. The cells were captured by wide-field microscopy using a color camera. Scale bars: 500 μm (N); 20 μm (A); 10 μm (B, D, E (nuclei), G, H, and O); 5 μm (nucleoli: C, I, and J); 3 μm (E, PML-NDS, and PML-NBs).

**Supplementary Figure 5. Inhibition of ATM, ATR, and RAD51 suppressed the formation of I-PpoI-induced PML-NDS.** (**A**) The efficiency of used concentrations of ATM and ATR inhibitors was assessed by analyzing the level of the serine 15 phosphorylated form of p53 (pS15-p53) by western blotting in RPE-$1^{hTERT}$ after doxorubicin treatment (0.56 µM). The inhibitors were added 1 h before doxorubicin. The quantification of the obtained signals is shown as $log_2$ of fold change (FC) between the ATM/ATR inhibition and control as a means from three biological replicates. (**B**) The I-PpoI was activated for 24 hours in RPE-$1^{hTERT}$-I-PpoI (clone 1A11). The number of 53BP1 (a marker of DSB) was assessed by indirect IF and ScanR analysis software after recovery from 24-hour-long activation of I-PpoI. The B02 and NU-7441 were added simultaneously with the activation of I-PpoI to restrict RAD51 filamentation or DNA PK. The data from three independent biological replicates (values for 200 nuclei) were pooled together and represented as histograms showing the frequency of nuclei (%) with the same number of 53BP1 foci. The bin center used for analysis was 2. (**C**) The number of nuclei with PNAs upon the conditions described in (B) was obtained by detection of PML using indirect IF and ScanR. The quantification was done manually by the evaluation of PML localization in more than 200 nuclei. (**D**) The efficiency of RAD51 knockdown was assessed 1 day after post-transfection with esiRAD51 by analyzing the level of RAD51 protein by western blotting in RPE-$1^{hTERT}$. The quantification of the obtained signals is shown as $log_2$ of FC between the esiRAD51 and pool of non-targeting siRNAs. Means from three biological replicates are shown. (**E**) The efficiency of LIG4 knockdown was assessed 1 day after post-transfection with esiLIG4 by analyzing the level of LIG4 mRNA by RT qPCR. The quantification of the obtained signals is shown as $log_2$ of FC between the esiLIG4 and pool of non-targeting siRNAs. Means from three biological replicates are shown. (**F)** RPE-$1^{hTERT}$-I-PpoI (1H4) were transfected by interfering RNA upon seeding. After 24 h, the I-PpoI was activated by doxycycline and Shield for another 24 h. Then, the medium was changed, and cells recovered from rDNA damage for 0 and 1 day. The control cells were treated with DMSO as MOCK at the same time. The γH2AX (a marker of DSBs; red) and PML (green) were detected by indirect IF and ScanR. The galleries of representative nuclei are shown. (**G**) RPE-$1^{hTERT}$-I-PpoI (1H4) were treated and analyzed as described in (F). The γH2AX (red) and PML (green) localization is shown after I-PpoI activation combined with KD of RAD51, LIG4, or B02. The nuclei and nucleoli were stained with DAPI (blue) and TOTO-3 (white), respectively. (**H** and **I**) RPE-$1^{hTERT}$-I-PpoI (1H4) were treated and analyzed as described in (F). The γH2AX count was obtained by ScanR analysis software. Then, the distribution analysis (%) was done in GraphPad (Prism) (see Figure 5G). The mean from four biological replicates shows the percentage of cells with γH2AX count higher than 35 foci (G) or between 0 – 2.5 foci (H). Scale bars: 20 µm (F and I). Results are presented as a mean ± s.d. Student's t-test was used for statistical evaluation. Asterisks indicate the following: ****$P < 0.0001$, ***$P < 0.001$, **$P < 0.01$, *$P < 0.05$

**Supplementary Figure 6. The resected DNA is present in G1 and S/G2 cells and colocalizes with I-PpoI-induced PNAs.** (**A**) The I-PpoI was activated 24 hours after RPE-1hTERT-I-PpoI (1H4) seeding using doxycycline and Shield. The cells were harvested 6, 8, and 24 hours after this, and RAD51(red) and PML (green) were detected by indirect IF and ScanR. The DNA was stained by DAPI (blue). The G1 and S/G2 cells were estimated according to the total DAPI fluorescence. The galleries of nuclei made from estimated gates (G1; S/G2) are shown. (**B**) The same experimental setup as described in (A) was used, but RPA2pS33 (red) and PML (green) were detected by indirect IF and ScanR. The DNA was stained by DAPI (blue). The G1 and S/G2 cells were estimated according to the total DAPI fluorescence. The galleries of nuclei made from estimated gates (G1; S/G2) are shown. (**C** and **D**) RPE-$1^{hTERT}$-I-PpoI (1H4) were transfected by interfering RNA upon seeding. After 24 hours, the I-PpoI was activated by doxycycline, and Shield (C) or DMSO was added (D) for another 24 hours. Then, the medium was changed, and cells recovered for 0 or 1 day. The PML and γH2AX were detected by indirect IF and ScanR. The scatter plots show the total DAPI fluorescence and mean γH2AX fluorescence estimated for each nucleus. The dashed red lines indicated the division to G1 and S/G2 subpopulations. (**F**) The presented columns indicate the percentage of nuclei with PNAs present in the G1 or S/G2 subpopulation after treatment described in C. Results are presented as a mean ± s.d. obtained from three biological replicates. Asterisks indicate the following: ****P < 0.0001, ***P < 0.001, **P < 0.01, *P < 0.05.
