## Supplementary Tables for "Topological stress triggers persistent DNA lesions in ribosomal DNA with ensuing formation of PML-nucleolar compartment"

Supplementary Table 1

| Class | Compound | Concentration | Specification | Citation |
|---|---|---|---|---|
| Topoisomerase inhibitors | camptothecin | 50 µM | TOP1 poison | (1,2) |
| | topotecan | 50 µM | TOP1 poison | (3) |
| | doxorubicin | 0.75 µM | TOP2 poison (< 1 µM), DNA intercalation, histone eviction | (4-6) |
| | aclarubicin | 0.05 µM | inhibition of TOP2 binding to DNA, DNA intercalation, histone eviction | (5,7) |
| | etoposide | 50 µM | TOP2 poison | (8,9) |
| RNAP I inhibitors | actinomycin D | 10 nM | DNA intercalation (preferentially to GC-rich sequences) | (10,11) |
| | BMH-21 | 0.5 µM | Intercalates DNA (preferentially to GC-rich sequences) | (12,13) |
| | CX-5461 | 5 µM | block the release of RNAPI pre-initiation complex | (14) |
| Inhibitors of rRNA processing | 5-FU | 200 µM | antimetabolite; inhibits pre-rRNA late processing | (15,16) |
| | MG-132 | 10 µM | The proteasome inhibitor; inhibits pre-rRNA late processing | (15,17) |
| | roscovitine | 20 µM | MAP-kinase inhibitor; inhibits pre-rRNA early processing | (15,18,19) |
| Replicative stress inducers | aphidicolin | 0.4 µM | DNA polymerase A, D inhibitor | (20) |
| | hydroxyurea | 100 µM | prevent dNTP accumulation at G1/S and block DNA synthesis | (21) |
| Others | oxaliplatin | 10 µM | alkylating agent, DNA cross-linking, in higher concentrations inhibition of RNA and protein synthesis | (22) (23) |
| | BrdU | 100 µM | nucleotide analogue; induces DNA damage response radio-sensitization | (24-26) |
| | IR | 10 Gy | DNA-damaging treatment | |
| | IFNγ | 5 ng/mL | JAK-STAT pathway activator; induces the expression of PML | (27,28) |

Supplementary Table 2

| | | | | |
|---|---|---|---|---|
| Antibodies used for immunofluorescence | | | | |
| anti-53BP1 | mouse, monoclonal | MAB3802 | Sigma-Aldrich/Merck (Darmstadt, Germany) | 1:400 |
| anti-BrdU | mouse, monoclonal | B8434 | Sigma-Aldrich/Merck (Darmstadt, Germany) | 1:500 |
| anti-DHX9 | rabbit, polyclonal | NB110-40579 | Bio-Techne/NovusBiological (Minneapolis, USA) | 1:400 |
| anti-PML | mouse, monoclonal | sc-966 | Santa Cruz Biotechnology (Dallas, TX, USA) | 1:100 |
| anti-PML | rabbit, polyclonal | sc-5621 | Santa Cruz Biotechnology (Dallas, TX, USA) | 1:200 |
| anti-PML | rabbit, polyclonal | ABD-030 | Jena Bioscience (Jena, Germany) | 1:500 |
| anti-phosphoserine 139 of histone H2AX | mouse, monoclonal | 05-636 | Millipore/Merck (Darmstadt, Germany) | 1:500 |
| anti-phosphoserine 139 of histone H2AX | rabbit, polyclonal | 11174 | Abcam (Cambridge, UK) | 1:500 |
| anti-nucleophosmin/B23 | mouse, monoclonal | 32-5200 | Invitrogen/Thermo Fisher Scientific (Waltham, MA, USA) | 1:200 |
| anti-PAF49 | rabbit, polyclonal | ab92428 | Abcam (Cambridge, UK) | 1:250 |
| anti-Rad51 | rabbit, polyclonal | | Gift from Pavel Janscak (University of Zurich, Switzerland) | 1:500 |
| anti-RPA194 (H-300) | rabbit, polyclonal | sc-28714 | Santa Cruz Biotechnology (Dallas, TX, USA) | 1:200 |
| anti-phospho-RPA2 (pSer33) | rabbit, polyclonal | NB100-544 | Bio-Techne/NovusBiological (Minneapolis, USA) | 1:500 |
| anti-UBTF | rabbit, polyclonal | HPA006385 | Sigma-Aldrich/Merck (Darmstadt, Germany) | 1:100 |
| Alexa Fluor 488 goat anti-mouse | goat anti-mouse | A-11029 | Invitrogene/Thermo Fisher Scientific (Waltham, MA, USA) | 1:1000 |
| Alexa Fluor 568 goat anti-mouse | goat anti-mouse | A-11031 | Invitrogene/Thermo Fisher Scientific (Waltham, MA, USA) | 1:1000 |
| Alexa Fluor 488 goat anti-rabbit | goat anti-rabbit | A-11034 | Invitrogene/Thermo Fisher Scientific (Waltham, MA, USA) | 1:1000 |
| Alexa Fluor 568 goat anti-rabbit | goat anti-rabbit | A-11036 | Invitrogene/Thermo Fisher Scientific (Waltham, MA, USA) | 1:1000 |
| Antibodies used for immuno-FISH | | | | |
| anti-53BP1 | mouse, monoclonal | MAB3802 | Millipore/Merck (Darmstadt, Germany) | 1:250 |
| anti-PML | Rabbit, polyclonal | sc-5621 | Santa Cruz Biotechnology (Dallas, TX, USA) | 1:400 |
| anti-nucleophosmin/B23 | mouse, monoclonal | 32-5200 | Invitrogen/Thermo Fisher Scientific (Waltham, MA, USA) | 1:300 |
| Alexa Fluor 405 goat anti-mouse | goat anti-mouse | A-31553 | Invitrogene/Thermo Fisher Scientific (Waltham, MA, USA) | 1:500 |
| Alexa Fluor 488 goat anti-rabbit | goat anti-rabbit | A-11034 | Invitrogene/Thermo Fisher Scientific (Waltham, MA, USA) | 1:500 |
| Anti-Digoxigenin-Rhodamine, Fab fragments | Sheep anti-digoxigenin | 11207750910 | Sigma-Aldrich/Merck (Darmstadt, Germany) | 1:15 |
| Antibodies used for immunoblotting | | | | |
| anti-GAPDH | mouse, monoclonal | GTX30666 | GeneTEX (Irvine, CA, USA) | 1:10 000 |
| anti-p53 | mouse, monoclonal | sc-126 | Santa Cruz Biotechnology (Dallas, TX, USA) | 1:500 |
| anti-p53 | mouse, monoclonal | ab1101 | Abcam (Cambridge, UK) | 1:1 000 |
| anti-topoisomerase I | rabbit, polyclonal | HPA019039 | Sigma-Aldrich/Merck (Darmstadt, Germany) | 1:1 000 |
| anti-topoisomerase I | rabbit, monoclonal | ab109374 | Abcam (Cambridge, UK) | 1:1 000 |
| anti-topoisomerase II α | mouse, monoclonal | sc-365916 | Santa Cruz Biotechnology (Dallas, TX, USA) | 1:1 000 |
| anti-topoisomerase II β | rabbit, polyclonal | sc-13059 | Santa Cruz Biotechnology (Dallas, TX, USA) | 1:1 000 |
| IgG-HRP goat anti-rabbit | goat anti-rabbit | 170-6515 | Bio-Rad (Hercules, CA, USA) | 1:10 000 |
| IgG-HRP goat anti-rabbit | goat anti-rabbit | #A6154 | Sigma-Aldrich/Merck (Darmstadt, Germany) | 1:10 000 |
| IgG-HRP goat anti-mouse | goat anti-mouse | 170-6516 | Bio-Rad (Hercules, CA, USA) | 1:10 000 |
| IgG-HRP goat anti-mouse | goat anti-mouse | #A9044 | Sigma-Aldrich/Merck (Darmstadt, Germany) | 1:10 000 |
| anti-phospho-p53 (pSer15) | mouse, monoclonal | #9286 | Cell signaling (Danvers, MA, USA) | 1:1 000 |
| anti-GAPDH | mouse, monoclonal | sc-47724 | Santa Cruz Biotechnology (Dallas, TX, USA) | 1:20 000 |
| anti-Rad51 | rabbit, polyclonal | sc-8349 | Santa Cruz Biotechnology (Dallas, TX, USA) | 1:500 |
| anti-β-tubulin | mouse, monoclonal | | Gift from Pavel Dráber (IMG CAS, Prague; Czech Republic) | 1:10000 |

Supplementary Table 3

| Name of plasmid | Description | Source |
|---|---|---|
| pCDNA4TO-FKBP-PPO-HA | Plasmid with a regulable expression of I-PpoI | (1); a kind gift of Libor Macurek (IMG CAS, Prague; Czech Republic) |
| pLVX-TET-ONE-puro | All-in-one lentiviral vector with puromycin resistance and strong inducible expression of genes using the Tet-On® system | Clontech Laboratories, Inc.; A Takara Bio Company |
| pCDH-CMV-MCS-EF1-Neo | Cloning and Expression Lentivector | System Biosciences, LLC (Palo Alto, CA 94303) |
| pLVX-TET-ONE-neo | All-in-one lentiviral vector with neomycin resistance and strong inducible expression of genes using the Tet-On® system. | This work |
| pLVX-TETOne-neo-FKBP-PPO-HA | lentiviral vector with neomycin resistance enabling the preparation of stable cell lines with a regulable expression of I-PpoI. | This work |
| pUC-hrDNA-12.0 | Plasmid bearing DNA of rDNA intergenic spacer. The source for the FISH probe targeting rDNA locus. | (2)a kind gift of Prof. Brian McStay (NUI Galway, Ireland) |
| BAC_CH507-535F5 | BAC bearing DNA of human distal junction. The source for the FISH probe targeting DJ locus. | BACPAC Resources Center (Emeryville, CA, USA) |


1. Warmerdam, D.O., van den Berg, J. and Medema, R.H. (2016) Breaks in the 45S rDNA Lead to Recombination-Mediated Loss of Repeats. *Cell reports*, **14**, 2519-2527.
2. van Sluis, M., van Vuuren, C., Mangan, H. and McStay, B. (2020) NORs on human acrocentric chromosome p-arms are active by default and can associate with nucleoli independently of rDNA. *Proc Natl Acad Sci U S A*, **117**, 10368-10377.


Supplementary Table 4

| Name of primer | Sequence |
|---|---|
| GA-PpoI-LVXpur-fr-F | ACTTCCTACCCTCGTAAAGAAGCTTGGTACCGAGCTCG |
| GA-PpoI-LVXpur-fr-R | GCAGGGGAGGTGGTCTGGATCCCCATAGAGCCCACCGCAT |
| GA-V-LVXpuro-F | GTTCTTCTGAACGCGTCTGGAACAATCA |
| GA-V-LVXpuro-R | GTTCAATCATGGTCGCGTTTTGCAAAAG |
| GA-F-neo-R | CCAGACGCGTTCAGAAGAACTCGTCAAGAAGGCGATAGAAG |
| GA-F-neo-F | AAACGCGACCATGATTGAACAAGATGGATTGCAC |
| LIG4_F | GAAGGCATCTGGTAAGCTCG |
| LIG4_R | AGCTTCCTTAGCACTCACACA |
| GAPDH fw | GTCGGAGTCAACGGATTTGG |
| GAPDH rev | AAAAGCAGCCCTGGTGACC |
